# Dynamics of odor-source localization: Insights from real-time odor plume recordings and head-motion tracking in freely moving mice

**DOI:** 10.1101/2023.11.10.566539

**Authors:** Mohammad F. Tariq, Scott C. Sterrett, Sidney Moore, Lane, David J. Perkel, David H. Gire

## Abstract

Animals navigating turbulent odor plumes exhibit a rich variety of behaviors, and employ efficient strategies to locate odor sources. A growing body of literature has started to probe this complex task of localizing airborne odor sources in walking mammals to further our understanding of neural encoding and decoding of naturalistic sensory stimuli. However, correlating the intermittent olfactory information with behavior has remained a long-standing challenge due to the stochastic nature of the odor stimulus. We recently reported a method to record real-time olfactory information available to freely moving mice during odor-guided navigation, hence overcoming that challenge. Here we combine our odor-recording method with head-motion tracking to establish correlations between plume encounters and head movements. We show that mice exhibit robust head-pitch motions in the 5-14Hz range during an odor-guided navigation task, and that these head motions are modulated by plume encounters. Furthermore, mice reduce their angles with respect to the source upon plume contact. Head motions may thus be an important part of the sensorimotor behavioral repertoire during naturalistic odor-source localization.

## Introduction

Animals exhibit rich sensorimotor transformations as part of their natural behavior. Studying naturalistic behaviors in rodents [1–3] offers the opportunity to understand the neural circuits involved in processing sensory information for goal-directed behaviors. For example, odor plume-guided navigation [4,5] is a tractable system in this pursuit, but has historically been under-studied due to the inability to experimentally monitor the complex odor plumes [6,7] that rodents encounter during free behavior. Lewis et al. [8] and Ackels et al. [9] have shown that the olfactory bulb of mice can encode the fast dynamics present in a naturalistic plume, but how this neural activity shapes behavior in response to intermittent plume encounters during naturalistic behaviors has not been clearly established.

Vickers and Baker [10] were the first to precisely correlate plume contacts to behavior in freely flying moths. Their seminal work, along with subsequent studies [11–16] revealed robust behavioral sequences exhibited by invertebrates during complex odor-source localization tasks. Additionally, walking flies navigating a turbulent odor plume also exhibit behavioral changes in response to plume encounters [17].

Mammalian responses to plume encounters during odor tracking using airborne plumes have not been characterized. Findley et al. [18] have recently shown head motions to be a key behavioral feature during an odor gradient-dependent search. However, the plume encounter-dependent changes in rodent head motions were not established in their study. In addition, Liao and Kleinfeld [19] have shown alterations in behavioral states of freely moving rats engaged in an exploratory behavior. They, along with Findley et al. [18], have also established important correlations between rodents’ sniffing and their head motions. We have previously used metal oxide sensors to record real-time olfactory information during airborne odor-tracking in mice [20]. Here we combine this method with head-mounted accelerometers to study how head motions change around plume encounters.

Like Findley et al. [18], and Liao and Kleinfeld [19] we establish head motions as a critical component of the mouse natural behavioral repertoire. We extend upon these findings by demonstrating that plume encounters in a turbulent setting modulate these head motions. Specifically, we show that preceding plume contacts mice increase the amplitude of their head-pitch motions in the active search band (5-14 Hz), with an immediate reduction after contact. In addition, mice lower their heads and switch to foraging after plume contacts, with a corresponding increase in the frequency of the head-pitch motions and decrease in the body angle with respect to the source. Our results thus suggest that mice are able to gather information about the relative orientation of the plume with each odor encounter.

Similar head motions are also involved in visually-guided tasks [21–23], highlighting the importance of head motions for sensory acquisition and motor guidance [3] during free behavior. Hence, our work combined with that of Findley et al. [18] establishes the groundwork to study the neural mechanisms underlying sensorimotor transformations during the naturalistic behavior of odor-plume tracking in mammals.

## Materials and Methods

### Animal Housing and Protocol

All animal procedures were approved by the University of Washington Institutional Animal Care and Use Committee (IACUC protocol #4356-01). C57BL/6J mice of both genders (n=7; 3 females/4 males), obtained from The Jackson Laboratory (Strain #000664) were used as test subjects. Mice were maintained on a reverse 12 hrs light schedule (7AM lights off; 7PM lights on). After surgery, described below, each mouse was housed individually.

### Surgery

All the mice were between 19-24 weeks of age at implantation. Each mouse underwent aseptic surgery using Isoflurane as anesthesia for implantation of a 3D printed part (S1 File), housing the female end of PCB connectors (Mill-Max 833-93-100-10-001000; 2 rows of 3 pins). The 3D part, printed using the Form2 3D-Printer (Formlabs) with tough resin and weighing 1g, was affixed to the skull with cyanoacrylate. A 1-mm screw was also implanted over the caudal skull on top of cerebellum to anchor the part. The glue was then built around the part and the screw to ensure long-term stability of the part. The exposed area of the skull was covered with dental cement. After each surgery, the mouse was monitored for 45 minutes in a bedding and chips free cage until the mouse was mobile again. The mouse was then housed individually and monitored for 10 days after surgery for post-surgery complications. Analgesics, Carprofen and topically applied Lidocaine, were administered before the surgery. Post-surgery carprofen was administered if needed.

### Behavioral training and task

After a >2weeks post-surgery period, the mice were water-restricted and trained to associate the smell of ethanol with water. This was achieved by placing weigh boats containing water and ethanol-soaked pads taped underneath in the home cage. Each mouse was given approximately 15 minutes to consume the water from the weigh boat. The behavioral apparatus and the task design for odor-source localization were as previously described [20,24]. Briefly, the odor port in the custom-designed arena [2m (l), 0.9m (w), 0.9m(h)] was placed randomly in an x-y location within a 1m(l) x 0.9m(w) area on the opposite end of the arena with respect to the home cage.

Ethanol was released by the odor port in a constant air stream from outside the arena, controlled by a valve. The valve gated the flow of the carrier air stream into an ethanol vial. The odorized air was then carried by tubing to the odor port located in the arena. A negative air pressure generated by placing a fan outside the arena then turbulently guided the odorized stream from the odor port. The goal for the mouse was to enter the arena from the home cage, located outside the arena but accessible through a gated passage, and locate the source of the odor port to receive reward (water droplet). A 1-s tone of 1 kHz signified the end of the trial.

After each trial was completed, the mouse was coaxed back into the housing cage located outside the arena, and the odor port was repositioned for the next trial. After each repositioning of the odor port, the odor valve was turned on for a few seconds before the commencement of the next trial. The valve was turned off only after the trial was completed, hence, the ethanol vapors were continuously released throughout the trial. The order in which the mice were tested was randomized for each day.

The arena was illuminated solely with infrared lights so that the mouse could not see the port but an overhead camera (Basler acA640-90uc) could record the animal’s position. In addition, white noise was continuously played in the room to mask any acoustic cues. To minimize strain by the cables, a linear actuator (Progressive Automation PA-15), with a slip-ring commutator mounted on a small plexiglass sheet, to minimize electrical noise, was installed in the arena. The actuator motion was manually controlled by an experimenter with a switch, located outside the arena, based on the location of the mouse within the arena. Each mouse was weighed before water restriction to establish the baseline weight and was then maintained at a weight >0.85 times the baseline weight. In addition to the behavioral task, the mice were supplemented with water following the same protocol as the ethanol-water association, described above, to maintain their body weight.

As shown previously, for single measurement locations the odor presence is patchy over time (see Fig 5 in [20]) in our arena. These measurements were done simultaneously with a photoionization detector (PID), the current gold standard for measuring odor molecules in the olfactory field [25], and the metal oxide sensors, head-mounted in our study, showing adequate correlations between the two sensors. The use of the metal oxide sensors for plume sampling has also been verified by other labs [26–28].

With the random placement of the odor-source for each trial (see above) and the motion of the animal, the location and time at which the ethanol molecules are close to the animal for sampling will spatiotemporally vary. Hence, the head-mounted sensor provided us with a read-out of the presence of the ethanol molecules within the sampling space of the animal. We thus defined plume encounters through offline analysis of the signal from the head-mounted sensor (see below).

### Head-mounted odor sensor and accelerometer

A Figaro TGS 2620 sensor, as described previously [20], mounted on a custom-designed Printed Circuit Board (PCB; OSH Park, Oregon US) was used to record the real-time olfactory information. Also mounted on the PCB was a 3-axis accelerometer (Analog Devices ADXL 335). The accelerometer and the ethanol sensor were respectively powered by 3.3V and 5V outputs from an Arduino (Adafruit), and shared a common ground. The PCBs were screwed onto a 3D printed part (S2 File) using miniature screws (McMaster-Carr#96817A704) and the part also contained male PCB pins to connect with the 3D printed part glued onto the mouse skull (Total weight of the 3D part and PCB with wires: ∼3.3 g). A 30 Gauge, 8 conductor cable (McMaster-Carr#3890N21) was soldered directly onto the PCB for signal conduction. The outer insulation jacket was removed from the cable to make it more flexible to facilitate mouse locomotion.

The data were digitized using a National Instruments NI6212 DAQ, and acquired and stored using LabView scripts. To acclimate the mice with the weight of the 3D printed part, a bare part without the PCBs mounted was connected to the head piece after the odor-localization behavioral task each day (Bare 3D part weight: ∼1.8 g). This allowed each animal to have an additional weight of 1.5 g to be carried on the head during the odor-localization task in the behavioral arena. In addition, before the odor-localization task in the arena, a smaller version of the apparatus was used to acclimatize the mice with the head-mounted sensors in a bedding-free mouse cage.

### Signal Analysis

All the time-series analysis was done prior to analyzing the position data to minimize the location bias. Furthermore, the training for DeepLabCut (DLC) [29,30], described below, was conducted by a researcher not involved in the data collection.

### Head-mounted Sensors

All analysis code was written in MATLAB R2019a (MathWorks). The ethanol sensor signal was low-pass filtered (5Hz cutoff) and then deconvolved using a difference-of-two-exponentials kernel with τ_Rise_ and τ_Decay_ values set at 0.02 and 10s, respectively [20]. For plume-event detection, the slope of the cumulative sum of the signal around all time points was calculated for 1-s segments before and after a specific time point. The onset of an event was defined as the time point when the ratio of the forward and reverse slopes crossed a threshold. The threshold was determined by pooling data from all of the odor-localization task sessions, and sessions involving the presentation of a plume without an animal running in the arena. This allowed us to minimize spurious detections that might result from the wind as temperature changes affect the signal over longer time scales. Events that were separated by greater than 5s were pooled for the group data.

The accelerometer readings were calibrated before and after running animals by placing the head piece in various orientations. The voltage recordings from the accelerometer were converted to units of g by zeroing and scaling appropriately. The recordings were then low pass filtered (20Hz cutoff), and the derivative of the signals was taken to convert the acceleration to jerk [31]. The resulting jerk signal was then used for time-frequency analysis via the *cwt* (Continuous Wavelet Transform) function. For determining when the head was raised, a 1-s moving median of the Z- and X-axes was calculated. The median signal for the X-axis was then subtracted from the Z-axis. The data from all the animal trials were then pooled and a three Gaussian model was then fit to the distribution using a Gaussian Mixture Model algorithm (MathWorks File Exchange; Matthew Roughan Commit Date: 2007). This analysis was based on the work of Liao and Kleinfeld [19]. To determine the dominant frequency of head-pitch motions, the band with the highest power within the active search (5-14 Hz) was used.

The real trajectories were defined as the time-window around the onset of events (described above), while random trajectories were obtained by selecting equal time-sized windows with the onset time point (t=0) randomly selected from the trial trajectory when the animal was within the arena. However, we did exclude 1-s windows on either side of real events for the selection of the onset time for the random trajectories. Hence, the onset of real events was possible within -5 to -1 and +1 to +5 s intervals of the random trajectories.

### Position Data

We used DeepLabCut (DLC) [29,30] to train a deep neural network for mouse position tracking. Trial videos were cropped to the size of the arena. The neural network was trained with videos from recorded trials without wire occlusion to reduce external artifacts. Our trained neural network exhibited a test error of 2.33 pixels, satisfying the reported standard for human level accuracy of 2.7 pixels [29]. We then developed a MATLAB (2022b) package to aggregate and analyze the position data alongside the existing accelerometer and ethanol sensor datasets. DLC output was used for angle calculations.

To calculate the body angle, the mid-point between the neck and body coordinates was determined. Then the angle between the vector connecting the mid-point with the body, and the vector connecting the mid-point to the port was defined as the body angle with respect to the port. The direction of the angle was taken as positive if the body to mid-point vector was clockwise to the vector connecting the mid-point to port, and negative when it was counter-clockwise. The head angle was defined as the angle between the vector connecting nose with neck, and the vector obtained by a 180° rotation of the vector connecting neck with body. Hence, an egocentric right head turn was thus considered negative while a left head turn was considered positive.

### Statistics

To minimize noise due to erroneous DLC labeling, any of the labels with a likelihood less than 0.9 was set to NaN, removing those points from the calculation. The angles were analyzed using the MATLAB circular statistics toolbox [32]. In addition to the time criteria (described above), only events greater than 15 pixels from the source, approximately the length of an adult mouse, at the time of encounters were pooled together for the group data (206 plume encounters across 111 trials for the accelerometer data). For the directional analyses, 281 plume encounters across 137 trials were grouped (additional trials with non-functional accelerometer were included into this data set).

Due to the low number of plume encounters across the trials and the inherent variability intrinsic to a naturalistic behavior, we took the timing of the plume encounter as the main driving variable for the behavioral changes. Hence, for statistical comparisons over time we took the mean of each trajectory’s parameter values within each 1s interval in the -5 to 5s time window and then compared the group distributions for the real and random trajectories within each 1s bin using Kolmogorov-Smirnov Test.

Table 1 shows the probability outcome for each 1-s bin for the different parameters. Since the random trajectories were taken from each trial, the relative contribution of each animal to the real and random distributions was similar. For comparing the distributions collapsed over time, we paired the strength of the resultant vector of the angles before the plume encounter with the strength of the resultant vector of the angles after the encounter for each trajectory and statistically compared them using the Paired Wilcoxon signed-rank test.

**Table 1.** Statistics Table.

| Comparison |  | Figure |  |  |  |
| --- | --- | --- | --- | --- | --- |
| Variable | Time | 2E | 3C | 3D | 5E |
| Real vs. Random | -5 to -4 | 0.6550 | 0.4722 | 0.3144 | 0.4801 |
|  | -4 to -3 | 0.8454 | 0.1310 | 0.2369 | 0.3794 |
|  | -3 to -2 | 0.8290 | 0.5793 | 0.4396 | 0.3002 |
|  | -2 to -1 | 0.1635 | 0.1362 | 0.2357 | 0.1867 |
|  | -1 to 0 | 0.0104 | 0.0007 | 0.1190 | 0.1154 |
|  | 0 to 1 | 0.0235 | 0.00002 | 0.0569 | 0.0292 |
|  | 1 to 2 | 0.0657 | 0.0456 | 0.0014 | 0.5007 |
|  | 2 to 3 | 0.0660 | 0.2673 | 0.1764 | 0.0890 |
|  | 3 to 4 | 0.0088 | 0.5906 | 0.0158 | 0.0091 |
|  | 4 to 5 | 0.0155 | 0.4567 | 0.0892 | 0.1866 |

Statistical probability comparing the real and random trajectories’ parameter values in the corresponding Figs within 1 s bins in a -5 to 5 s window around plume encounters. Significant bins are shaded gray with the significance level indicated by the different shades of gray.

## Results

To understand the plume-encounter dependent changes in mouse head motions during airborne-odor source localization, we combined our previously published method of recording real-time plume information [20] with head-motion monitoring using a 3-axis accelerometer.

### Concurrent head-motion tracking with real-time plume information during odor-guided navigation task

Fig 1A shows the 3D part designed to carry the Printed Circuit Board (PCB) with the ethanol sensor and accelerometer. The orientation of the accelerometer with respect to the ground can also be seen in Fig 1A. This device allowed us to record the real-time plume information experienced by, and head motions of a mouse engaged in odor-guided navigation.

An example trial showing the spatial trajectory of the mouse is presented in Fig 1B, while the accelerometer reading, speed, and olfactory information experienced by the mouse during that trial are presented in Fig 1C. To better visualize the evolution of the signals over time and space, zoomed-in views around the encounters labeled in Figs 1B and C are presented in Fig 1D. Hence, this method allowed us to examine how the head motions of mice engaged in the odor-localization task change with each odor-plume encounter.

**Fig 1.**
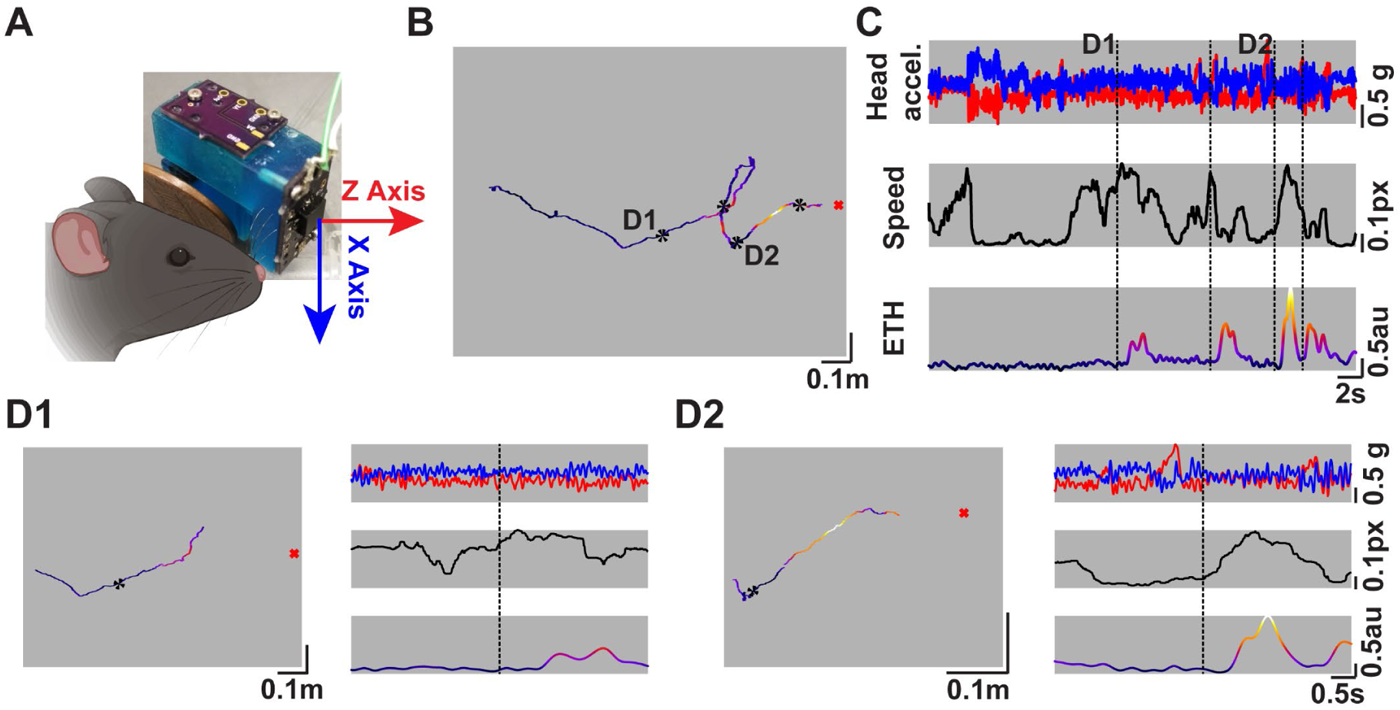
Concurrent head-motion tracking with real-time odor plume monitoring. **A)** The designed 3D-printed part with the custom designed PCB to monitor the plume information along with an accelerometer. The arrows correspond to the axis of orientation of the accelerometer and the color correspond with panel **C. B)** An individual trajectory of a mouse engaged in plume-tracking to locate the odor source for a water reward. Overlaid on the trajectory is the colormap of the real-time plume signal experienced by the mouse. The color-scale goes from dark (low signal) to lighter (higher signal) and corresponds with the colors shown in **C**. The black asterisks represent the locations of plume encounters. The red “x” marks the location of the odor source. **C)** (Top) The read-out over time from the Z- and X-axis channels of the accelerometer (red & blue, respectively) from the trajectory shown in **B**. (Middle) The speed of the animal over time during the trial shown in **B.** (Bottom) The plume signal experienced by the mouse during its search. The color-scale, same as **B**, goes from dark (low) to light (high). Black dashed lines represent the times of plume encounters. **D1)** A 3-s (before and after) window showing the position (left) and time (right) evolution of the signals around the encounter labeled as **D1** in **B** and **C**. **D2)** Same as **D1** but for the encounter labeled as **D2** in panels **B** and **C**.

### Plume encounter-dependent changes in the amplitude of the mouse head-pitch motions

To better characterize how plume encounters affect the head-pitch motions, we calculated the derivative of the X-axis signal from the accelerometer to convert the acceleration to jerk [31]. Since the X-axis is aligned with the direction of gravity, this channel was the most useful for studying the up and down (pitch) motions of the mouse head. It was readily apparent that the pitch motion had many frequency components embedded in the signal, as previously reported [18,19,31]. We therefore aligned the jerk signals with respect to the onset of plume encounters shown in Fig 2C, and conducted time-frequency analysis using the CWT of the jerk signal as shown in Fig 2D. The dominant frequency band was found to be in the 5-14 Hz range, the frequency band correlated with active exploration in freely moving rodents [19,33]. Interestingly, this frequency band was also prominent during the non-plume encounter portions as seen in the randomly selected trajectories (middle panels; Figs 2C-E).

To better characterize how the amplitude of head-pitch motions evolve over time, and in particular around plume encounters, we examined the power within the active search band (5-14 Hz) over time presented in Fig 2E. Additionally, the power from the example trial presented in Figs 1B-D is presented in Fig 2B. As seen in Fig 2E, there is a rapid increase in the amplitude of pitch motions immediately before plume encounters, which decreases rapidly soon after plume encounters. Thus, mice move their heads up and down more vigorously before plume encounters, and dampen these motions following an encounter.

**Fig 2.**
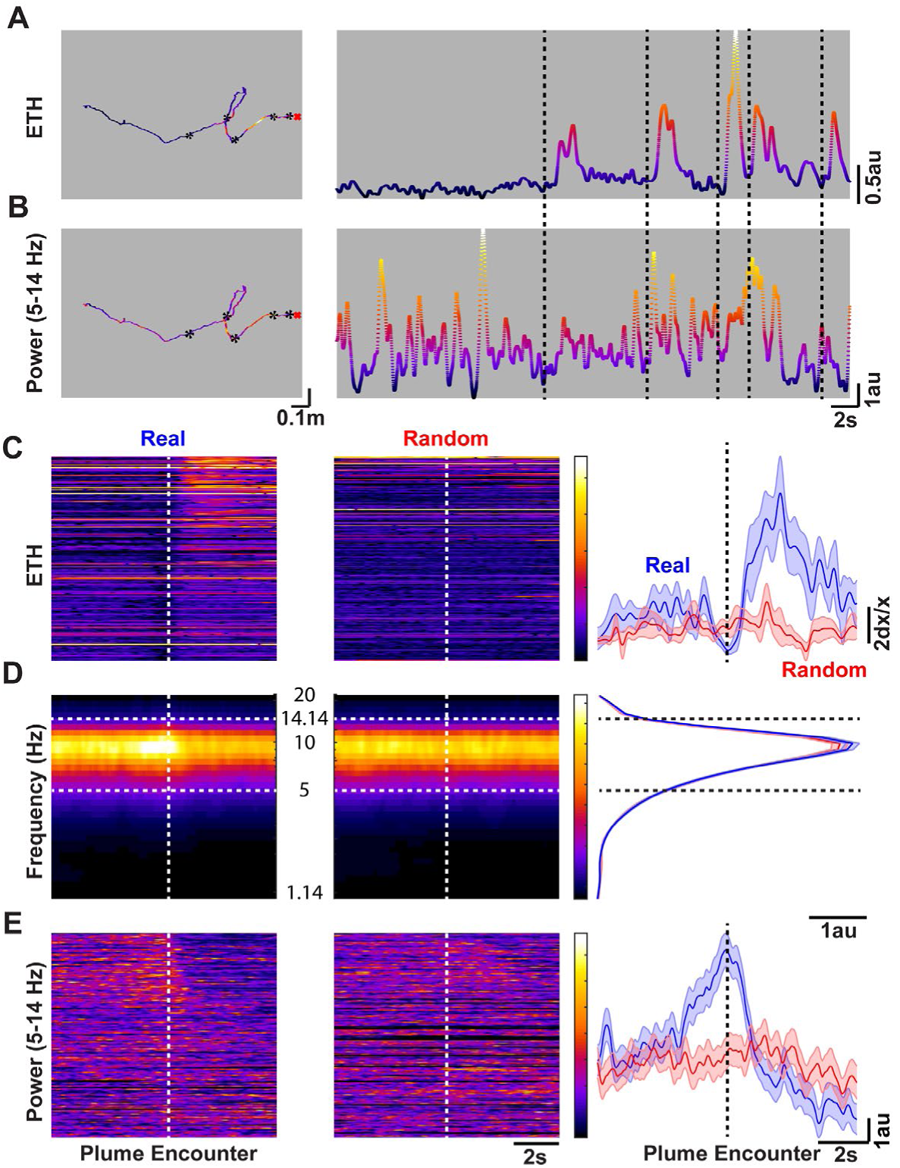
Plume encounters are preceded by increased amplitude of head-pitch motion in the active search (5-14 Hz) band, and immediately followed by decreased head-pitch motion. **A)** Same trajectory with the plume signal overlaid over space (left), and over time (right) as depicted in Figs **1B-D**. **B)** Amplitude of the mouse head-pitch motions in the 5-14 Hz band (see panel **D**) during the search shown in **A** over location (left) and time (right). The black asterisks, and dashed lines respectively represent the locations and times when plume encounters occurred. **C)** Group data showing the heat map of the plume signal in a 5-s window before and after a plume encounter for real (left) and randomly selected portions (middle), while the mean±SEM of all the encounters’ and random trajectories’ plume signals (Real: blue; Random: red) over time is presented (right). The signals in the left and middle heatmaps are sorted by the distance of the animal from the source when the encounter happened (Top: closest). **D)** A 5-s window (before and after plume encounter) CWT of the derivative of the accelerometer pitch motion showing 5-14 Hz predominant band (demarcated by the white dashed lines), referred to as the active search band, during real (left), and randomly selected trajectories (middle). The color scales are the same for the spectrograms for real and randomly selected data. In addition, mean power over the frequency bands are presented (right). Black dashed lines again demarcate the 5-14Hz band. **E)** The time-evolution of the power in the active search band during real (left) and randomly selected (middle) trajectories. The sorting of the signal is the same as in **C**. Furthermore, the color scale is the same for both heatmaps. Mean±SEM (right) of the power in the active search band over time is also presented.

### Mice lower their heads following plume encounters

Liao and Kleinfeld [19] have shown that rats engaged in free exploration switch from rearing to foraging, and that the sniff frequency during the foraging and rearing epochs are different. To test if such switching also occurred during our task, we examined the difference between the moving medians of the Z- and X-axes. This is an indirect method to look at the relative angle of the head with respect to the ground.

The distribution of this difference across all animals is presented in Fig 3A. This distribution was then fit to a sum of three Gaussians (black dashed curve) and the third Gaussian (magenta) was then used to establish the threshold for raising of the head [19]. To determine the number of Gaussians, we applied the Bayesian Information Criteria (BIC). The BIC decreased substantially with three Gaussians, and marginally decreased with higher numbers so we chose the three Gaussian model. We next looked at the dominant frequency of head-pitch motions during epochs when the head was either raised (magenta) or lowered (cyan) as shown in Fig 3B. The leftward shift of the dominant frequency during head-raising is consistent with that observed by Liao and Kleinfeld [19].

Furthermore, to study how plume encounters affected the position of the head we looked at this difference measure over time for the trajectories presented in Figs 2C-E. As shown in Fig 3C, there is an increase followed by a decrease right around plume encounters, suggesting that the mice stop scanning the air upon plume contacts. This is also accompanied with a corresponding increase in the dominant frequency of the head pitch motion as shown in Fig 3D. While we cannot correlate these head pitch-motion changes with changes of sniffing frequency in our paradigm, many groups [18,19,31] have established a strong correlation between sniffing and rodent head pitch motions in this frequency band.

**Fig 3.**
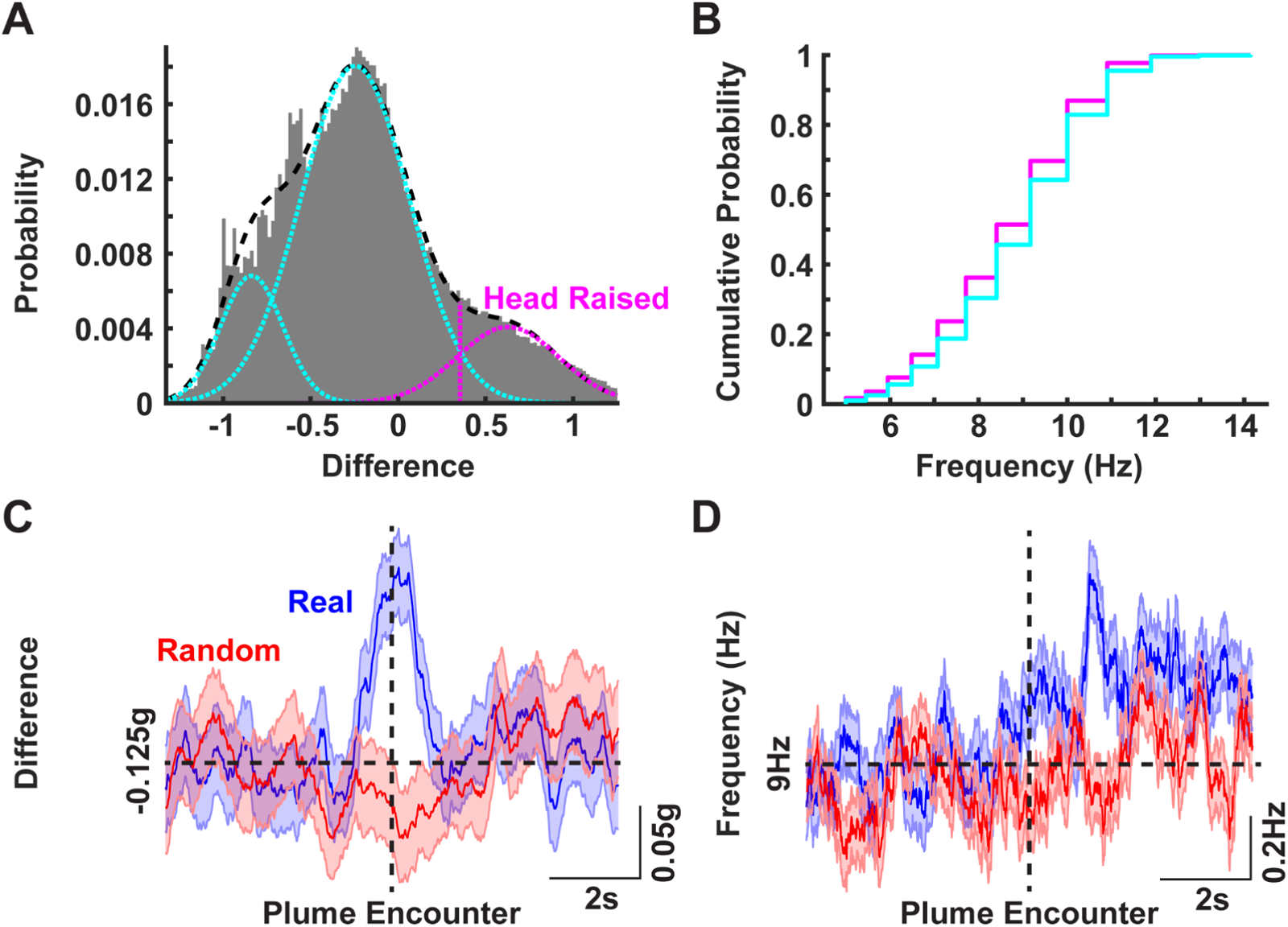
Plume encounter results in a lowering of the head with a corresponding increase in the frequency of head-pitch motion. **A)** The difference in the median Z- and X-axes read out from the accelerometer pooled over all trials and all animals. The resulting distribution was then fit with the sum of three Gaussians (black dashed curve). The mean-S.D. (magenta dashed line) of the third Gaussian (magenta curve) was then set as a threshold to classify the head as either raised (values greater than the dashed magenta line) or lowered. **B)** Cumulative histograms of the dominant frequency of head-pitch motion in the active search band during the head-raised (magenta) and head-lowered (cyan) epochs. A leftward shift in the pitch-motion frequency is apparent during head-raised epochs. **C)** Mean ± SEM of the difference of median of Z- and X-axes readings for real (blue) and randomly selected (red) trajectories (same plume encounter events as those presented in Figs **2C-E**). **D)** Mean ± SEM of the dominant frequency of the head-pitch motions around plume encounters for real and randomly selected trajectories.

### Mice reduce their body angle with respect to the source upon plume encounters

We next asked if mice can determine the relative location of the source with each plume encounter. To answer this question, we calculated the body angle with respect to the source (see Methods). Fig 4A shows the body angles of the mouse over time for the trajectories presented in Fig 1D. It was apparent that each mouse reduced the variability of its angle with respect to the source after plume encounter. To further quantify this effect, we calculated the mean angle of each trajectory 3s before and after the encounter, and then calculated the difference between the instantaneous angles and the mean angles for that time period. Fig 4B shows the probability heatmap over time for this angular difference for the real (blue; top) and random (red; bottom) trajectories. The distribution of the difference angles for the real trajectories becomes narrower (blue) after plume encounters than before encounters (gray), as can be seen in Fig 4C. No such change is observed for the randomly selected trajectories (Fig 4C; p = 0.0002 real & p = 0.6383 randomly selected trajectories; Paired Wilcoxon signed-rank test).

Furthermore, the mean angle over time reduces after plume encounter, as can be seen in Fig 4D with a corresponding increase in the strength of the resultant vector for real (blue) trajectories. The mean angle for the randomly selected trajectories (red; Fig 4D top), on the other hand, hovers around 10° (close to no particular direction preference with respect to the source). Furthermore, the high vector strength for the random trajectories (red; Fig 4D bottom) signifies increased proportions of angles randomly distributed within a range of ±90° around the mean angle.

**Fig 4.**
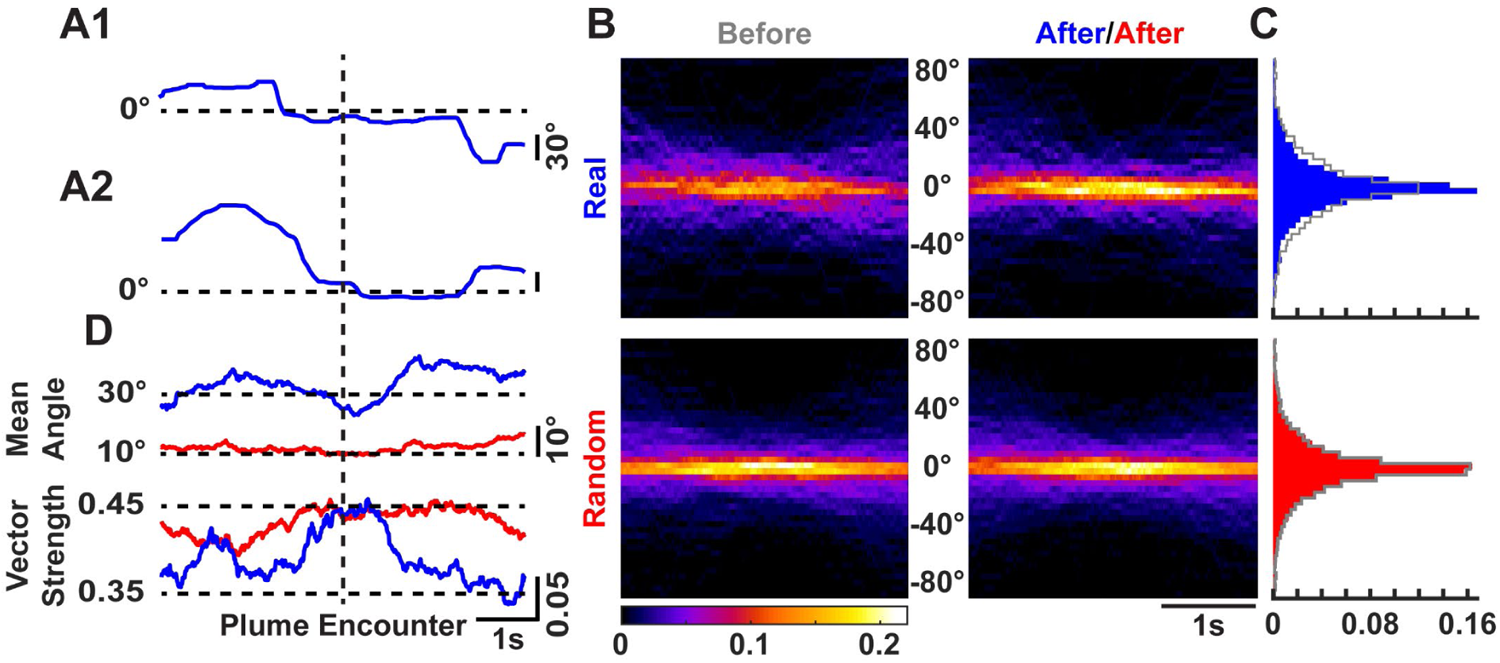
Mice reduce their body angle with respect to the odor source upon plume encounter. **A1)** The body angle (see Methods for description) of the mouse with respect to the odor source during the trajectory **D1** shown in Fig 1. **A2)** Same as **A1** but for the trajectory **D2** in Fig 1. **B)** The probability heatmap of the angle difference between instantaneous angles and the mean angle 3s before (gray) and after for each real (blue; top) and randomly selected (red; bottom) trajectory over time. **C)** The distribution of the difference in instantaneous angles and the mean angles before (gray) and after plume encounter for real (blue; top) and randomly selected (red; bottom) trajectories. Notice no difference between the before and after distribution for randomly selected trajectories (red; bottom), while the distribution after encounter for real trajectories (blue; top) becomes narrower than the distribution before (gray; top) plume encounter. **D)** The mean angle (top) over time for real (blue) and randomly selected (red) trajectories with the corresponding strength (bottom) of the resultant vector.

### Plume encounters reduce mouse head yaw angle variability

Casting crosswind of an odor source has been established as an efficient strategy to locate the plume in the horizontal plane for flying insects [10,12,13]. We therefore looked at the head yaw angle as a correlate for an active sensing strategy for walking mice to locate the plume in the horizontal plane. Fig 5A (top) shows the mean angle over time for the real (blue) and randomly selected (red) trajectories. While the mean angle for the real and randomly selected trajectories do not differ after plume encounters, there is a slight increase in the strength of the resultant vector (bottom) for real trajectories as compared to the random. This increase in strength for the real trajectories suggests that variability around the mean angle decreases within a 1s period before and after the plume encounters.

**Fig 5.**
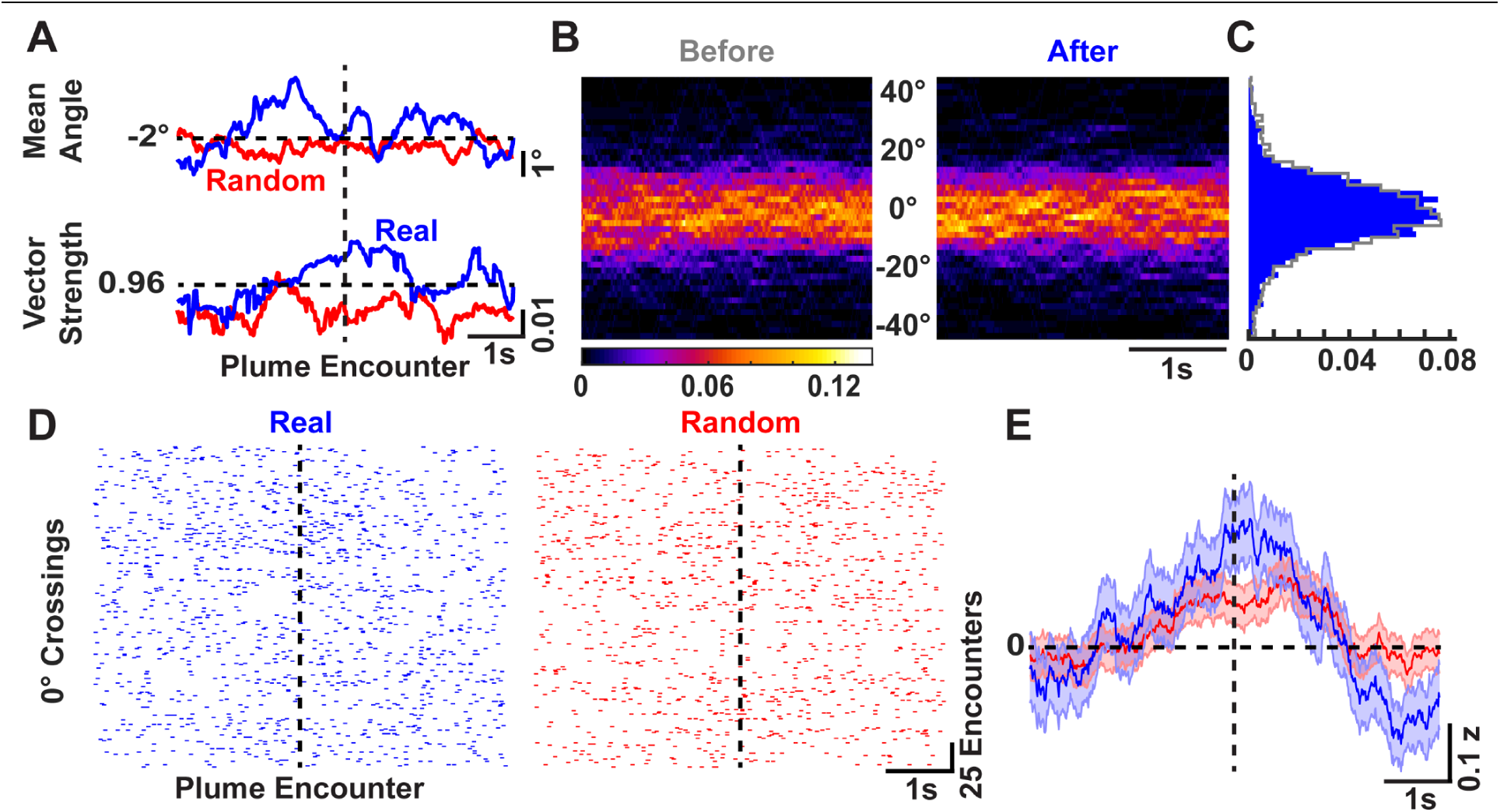
Mice reduce the variability of the yaw angle of head motions after plume encounter during odor-guided search. **A)** The mean angle (top) over time for real (blue) and randomly selected (red) trajectories with the corresponding strength of the resultant vector (bottom). Notice the increase in the strength peri-plume encounter for the real trajectories. **B)** The probability heatmap over time for left (+ve) and right (-ve) head motions before (grey) and after (blue) plume encounter. **C)** The head angle distribution before (grey) and after (blue) plume encounters. **D)** Raster plots for 0° crossing events for real (blue; left) and randomly selected (red; right) trajectories. **E)** Mean ± SEM of the z-scored 0° crossing rolling rates for the real (blue) and random (red) trajectories.

In addition, the probability heatmap over time before (grey) and after (blue) plume encounters is presented in Fig 5B, with the distribution of the yaw angles presented in Fig 5C. The distributions collapsed over time did not show a significant difference for real and random trajectories (p = 0.2036 for real & p = 0.5980 for random trajectories; Paired Wilcoxon signed-rank test). To further test if this reduction in the yaw angle variability around the plume encounters was due to increased rate of yaw turns, we looked at the number of 0° crossings before and after plume encounters. This measure is presented as raster plots for the real (blue) and random (red) trajectories in Fig 5D, where each tick represents a 0° crossing event. Fig 5E shows the mean ± SEM of the z-scored rolling rate of the 0° crossings for the real (blue) and random (red) trajectories. There is a transient increase in the rate for the real trajectories right after plume encounter while a decrease in the rate later after the encounter.

## Discussion

Correlating sporadic plume encounters to behavioral changes, and neural recordings during odor-guided navigation has remained an elusive goal [6,7] due to the constraints in real-time olfactory monitoring. Odor plume features have been shown to be encoded by the mouse olfactory bulb and guide behavior [8,9,34]. We previously reported a method to record the odor information available to a freely moving mouse using head-mounted sensors [20] that has been adopted by other groups to fit their research goals [28,35]. While these sensors can detect onsets of plume, we believe that more research in the sensor technology by engineers and chemists will also greatly benefit olfactory neuroscientists [36]. Here we combined our odor-sensing method with head-motion tracking using accelerometers to study how plume encounters shape head motions during an odor-localization task.

We show that mice actively search for olfactory plume encounters in our odor-source localization task by exhibiting robust head-pitch motions in the 5-14Hz range. These head motions are increased in their amplitude just before the plume encounter, and robustly decrease right after the encounter. This, in conjunction with our previously published result of a decrease in speed after plume encounters [20] suggests that the decrease in the head-pitch motion might be a strategy to localize the odor source. Interestingly, this reduction in ground speed right upon attractive odor presentation has also been recently shown in walking flies [37], and in freely flying flies upon optogenetic activation of the olfactory receptors [38].

Movement has been shown to generate and modulate neural signals in sensory brain areas possibly to account for the movement of the sensors [3,39–42]. Hence, studying how and if the olfactory neural system accounts for these robust head movements during odor-guided navigation is potentially a fruitful future research avenue. The vigorous head motions before encounter could be made to catch a whiff of the plume, while the reduction in the amplitude with a concomitant increase in the frequency of the head-pitch motions could be useful to more precisely decipher the location of the source. Our findings complement a recent publication that shows increase in the respiration frequency of freely moving mice during exploration of food odor, and novel object and odor [43].

Consistent with this two-stage model for source localization, we find that following plume encounters mice reduce their angle of travel with respect to the source. This suggests that mice can extract meaningful information about the relative orientation of the source, or the centerline of the plume from each plume encounter. Recent work in walking flies using optogenetic activation of olfactory receptors in a plume-like manner has shown that flies can sense the direction of the motion of odor molecules even when there is a mismatch between the wind direction and the odor motion [44].

Our results of the reduction of the angle with respect to the source along with a decrease in the variability of the angles then complements the odor-motion sensing study in flies. We also show that mice lower their heads after plume encounters, with an increase in the frequency of their head motions. This lowering of the head is consistent with normative theoretical strategies of switching to sampling the air vs. sampling closer to the ground for a computational searcher [45]. While we did not test for the sex-specific differences, a recent study [46] has shown that male and female rats differ in sampling durations of certain odors and odor-induced electrophysiological changes in the olfactory bulb. Studying plume encounter-dependent changes between sexes in a bigger cohort of animals would be an interesting next step.

In addition, we show that the yaw head angles are reduced in their variability after plume encounters, with a subtle increase in the rate of yaw turns again supporting the idea that mice change sampling strategy after a plume encounter [47]. While many mammals, including humans, have been shown to adapt their sensing strategies while following odor trails [18,48–50], here we show that mice adapt their strategies locked to plume encounters during odor-guided navigation. Furthermore, the evolution of the change in the different behavioral parameters suggest switching of behavioral motifs [51].

Together, our observations support a model in which mice engage in search behavior prior to plume contact, then adjust this behavior upon a plume encounter probably to extract meaningful information about the relative orientation of the source. While previous models of plume-guided search behavior emphasize the need for repeated plume sampling over space and time to iteratively estimate source position, the results we present instead suggest that mice have evolved a robust behavioral strategy, which combined with morphological and neural adaptations, may enable them to extract useful directional cues from even single plume encounters. Our results suggest that mice, and perhaps all macrosmatic mammals, receive and process vastly more directional information from odor plumes than previously thought.

## Acknowledgements

We like to acknowledge the members of the Perkel, and the Gire labs for helpful comments and discussions on this project.

## Author Contributions

**Conceptualization:** M.F.T., D.J.P., D.H.G.

**Data Curation:** M.F.T., L.

**Formal analysis:** M.F.T., S.C.S., L.

**Funding acquisition:** D.H.G.

**Investigation:** M.F.T., S.M.

**Writing – original draft:** M.F.T.

**Writing – review & editing:** M.F.T., S.C.S., L., D.J.P., D.H.G.

